# Bacterial plasma membrane is the main cellular target of silver nanoparticles in *Escherichia coli* and *Pseudomonas aeruginosa*

**DOI:** 10.1101/322727

**Authors:** Olesja M Bondarenko, Mariliis Sihtmäe, Julia Kuzmičiova, Lina Ragelienė, Anne Kahru, Rimantas Daugelavičius

## Abstract

Silver nanoparticles (AgNP) are widely used in consumer products, mostly due to their excellent antimicrobial properties. One of the well-established antibacterial mechanisms of AgNP is their efficient contact with bacteria and dissolution on cell membranes. To our knowledge, the primary mechanism of cell wall damage and the event(s) initiating bactericidal action of AgNP are not yet elucidated.

In this study we used a combination of different assays to reveal the effect of AgNP on i) bacterial envelope in general, ii) outer membrane (OM) and iii) on plasma membrane (PM). We showed that bacterial PM was the main target of AgNP in Gram-negative bacteria *Escherichia coli* and *Pseudomonas aeruginosa*. AgNP depolarized bacterial PM, induced the leakage of the intracellular K^+^, inhibited respiration and caused the depletion of the intracellular ATP. In contrast, AgNP had no significant effect on the bacterial OM. Most of the adverse effects on bacterial envelope and PM occurred within the seconds and in the concentration range of 7-160 μM AgNP, depending on the bacteria and assay used, while irreversible inhibition of bacterial growth (minimal bactericidal concentration after 1-h exposure of bacteria to AgNP) occurred at 40 μM AgNP for *P. aeruginosa* and at 320 μM AgNP for *E. coli*.

Flow cytometry analysis showed that AgNP were binding to *P. aeruginosa* but not to *E. coli* cells and were found inside the *P. aeruginosa* cells. Taking into account that AgNP did not damage OM, we speculate that AgNP entered *P. aeruginosa via* specific mechanism, e.g., transport through porins.

## INTRODUCTION

Silver nanoparticles (AgNP, 1-100 nm-sized) are most widely used NP in consumer products, mostly due to their excellent antimicrobial properties (1). However, the mechanisms of antibacterial action of AgNP are still under debate. Recent comprehensive review summarized four main proposed mechanisms of NP-bacteria interaction explaining the antibacterial effects of metal oxide NP including (i) generation of reactive oxygen species (ROS), (ii) partial dissolution and release of toxic metal ions, (iii) NP accumulation on membrane surface and (iv) internalization of NP (2). Although all these mechanisms have been shown to apply also in case of AgNP, the prevailing mechanism of toxic action of antibacterial properties of AgNP is the oxidative dissolution on bacterial surface (3). Indeed, in our previous study we showed that the interaction of NP with bacterial cell wall accompanied by the release of toxic Ag^+^ ions that contributed significantly to the antibacterial mechanisms of various AgNP to different Gram-negative and Gram-positive bacteria (4). In addition, we observed that medically important pathogenic bacteria *Pseudomonas aeruginosa* were inherently more susceptible to AgNP (especially to the colloidal form of AgNP, Collargol) than *Escherichia coli* cells. The current study is a continuation of our previous work that aims to provide the detailed insights into the interactions of AgNP with Gram-negative bacteria *E. coli* and *P. aeruginosa* with the focus on the effects on bacterial cell membranes. We selected the colloidal form of AgNP, Collargol, as the model AgNP since it has a long history of usage and is prescribed as a medication in several countries (1).

Opportunistic pathogens *E. coli* and *P. aeruginosa* were selected as bacterial models. Both are Gram-negative bacteria, i.e., have two membranes in their cell envelope - the outer membrane (OM) and the plasma membrane (PM) that possess different composition and therefore, distinct functions (Fig. 1). OM is covered by the dense layer of highly negatively charged lipopolysaccharides (LPS) and serves as a selective permeability barrier. OM hinders the entrance of hydrophobic substances and macromolecules (5) but allows the entrance of small charged molecules into the periplasm through the porins - narrow channels for ions and nutrients. In contrast, PM is relatively permeable for hydrophobic compounds but not for the inorganic ions and hydrophilic compounds (6, 7). PM is of utmost importance for bacteria since all the membrane-related functions of prokaryotic cells are performed in the PM (e.g., respiratory chain functions, lipid biosynthesis, protein secretion and transport events) (5, 8).

**FIG 1.**
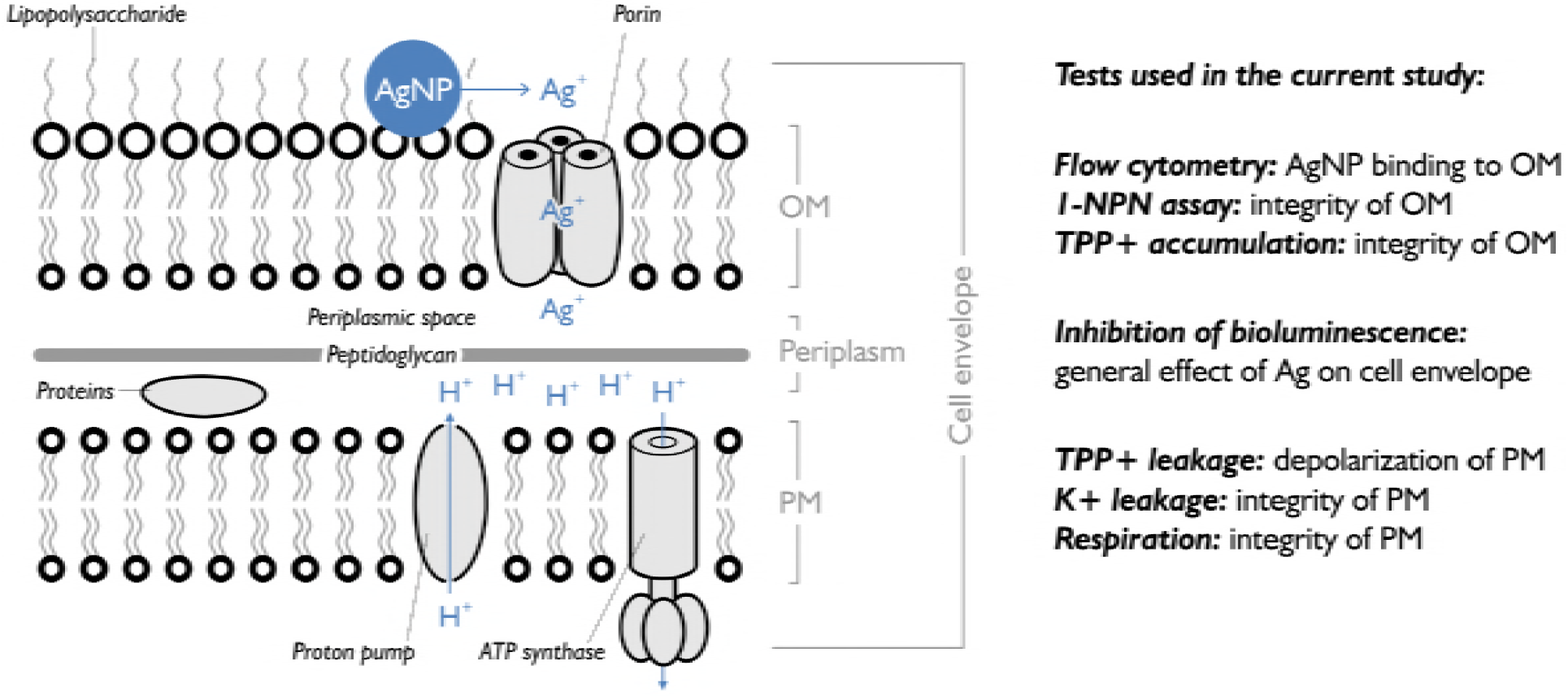
**Left:** Schematic structure of cell wall of Gram-negative bacteria, possible localization and transport mechanisms of Ag^+^ ions and Ag nanoparticles (AgNP). **Right:** Summary of the assays used in the current study and addressing the effects of studied compunds on cell envelope in general, bacterial outer mambrane (OM) or (plasma membrane) PM. 1-NPN, 1-N-phenylnaphthylamine; TPP^+^, tetraphenylphosphonium.

To reach the PM of Gram-negative bacteria, Ag ion/AgNP should first pass OM. Most likely, Ag^+^ ions traverse the OM through porins (9, 10, 11). Next, Ag^+^ ions induce several successive toxic events targeting PM. For instance, Dibrov et al (12) showed that Ag^+^ ions induce a proton (H+) leakage through the membrane vesicles of *Vibrio cholerae*. Authors suggested that H+ leakage is the main mechanism behind the bactericidal action of Ag^+^ ions in *V. cholerae*, and assumed that it occurs due to the unspecific, Ag^+^-induced modification of proteins or phospholipid bilayer.

In contrast to Ag^+^ ions, the mechanism by which AgNP interact with the cells and possess the antibacterial effect is not yet convincingly revealed. It is hypothesised that AgNP enter the cells by creating irregular-shaped pits on bacterial surface, inducing the LPS release and damaging the bacterial OM (10, 13, 14, 15). These events allow the penetration on AgNP into periplasm where AgNP act similarly to Ag^+^ ions, i.e., destroy functions of SH-groups containing enzymes due to the very high affinity of Ag ions towards sulfur (15).

The main aim of this study was to investigate the effects of Ag ions (AgNO_3_) and AgNP (Collargol, protein coated AgNP) on bacterial outer and plasma membranes in more detail. Effects of silver compounds on bacterial membranes were studied using different approaches: (i) **bioluminescence inhibition assay** to estimate the general damage of bacterial cell envelope and associated loss of energy (ATP) production; (ii) **bacterial viability testing** to determine minimal bactericidal concentration (MBC) of Ag compunds; (iii) **flow cytometry** to assess the bacteria-NP interaction and possible uptake of AgNP; (iv) **1-N-phenylnaphthylamine (1-NPN) assay** to estimate the integrity of OM, (v) **tetraphenylphosphonium (TPP^+^) assay** to discriminate the effect of Ag compounds on PM and OM and (vi) **K^+^ content and respiration** of cells to show the effects of Ag compounds on PM. All the assays addressed the short-term effects of studied compunds on cell envelope in general, bacterial OM or PM as depicted in Fig 1.

## MATERIALS AND METHODS

### Chemicals

All the chemicals were at least of analytical grade. Polymyxin B (PMB) was purchased from Sigma-Aldrich, 3,5-dichlorophenol (3,5-DCP) from Pestanal, AgNO_3_ from J.T.Baker, protein (casein)-coated colloidal AgNP from Laboratorios Argenol S. L. (batch No 297), propidium iodide from Fluka, tetraphenylphosphonium chloride (TPP^+^) from Sigma-Aldrich. The stock solutions of AgNO_3_ and AgNP were prepared at 2 mM (215.7 mg Ag/l, 20 ml) in autoclaved deionized (DI) water and further stored in the dark at +4 °C. The stock solutions were vortexed before each use. If not stated otherwise, all subsequent dilutions and experiments were performed in Ag^+^ ions non-complexing buffer consisting of 50 mM 3-(N-morpholino)propanesulfonic acid (MOPS) adjusted to pH 8 with (hydroxymethyl)aminomethane (Tris) and further referred as MOPS-Tris buffer.

### Characterization of silver nanoparticles

The shape, primary size and coating (casein) content of AgNP were determined in our previous publication (16). Hydrodynamic size (measured by dynamic light scattering, DLS), polydispersity index (pdi) and ζ-potential in MOPS-Tris buffer were measured at concentration 200 mg/l using Malvern Zetasizer (Nano-ZS, Malvern Instruments, UK). Dissolution of AgNP was quantified using atomic absorption spectroscopy (AAS). For that AgNP were incubated in MOPS-Tris at concentration 10 mg/l for 10 min, 1 h or 24 h, ultracentrifuged (390,000 g, 30-60 min) and the supernatant was analyzed by GF-AAS in accredited (ISO/IEC, 2017 (17)) laboratory of the Institute of Chemistry of Tallinn University of Technology, Estonia. The ratio of determined dissolved Ag to total nominal metal in AgNP was designated as dissolution (%).

### Bacterial strains and cultivation of bacteria

*E. coli* MC1061 (obtained from Prof. Matti Karp, Turku, Finland) and *P. aeruginosa* DS10-129 (isolated from soil, (18)) were used in all experiments. In addition, *Pseudomonas aeruginosa* PAO1 (obtained from Prof. Patrick Plesiat, Besanc, France) was used to confirm the results of TPP^+^, K^+^ and respiration assays. All cultivations were performed on a shaker at 200 rpm, 30°C. Bacteria were pre-grown overnight in 3 ml of NaCl-free Luria-Bertani medium (LB, 10 g tryptone and 5 g yeast extract per litre, pH=7). In case of bioluminescent *E. coli* and *P. aeruginosa* (see the next section), medium was supplemented with appropriate antibiotics (100 mg ampicillin per litre for *E. coli* and 10 mg tetracycline - for *P. aeruginosa*). 10-50 ml of fresh medium was inoculated with 1/20 diluted overnight culture and bacteria were grown until mid-exponential phase (OD_600_ of 1), centrifuged at 6000 g for 5 min, washed twice with equal amount of 50 mM MOPS-Tris and adjusted to OD_600_~1 corresponding to approximately 10^9^ cells/ml.

ζ-potential of bacteria was measured in MOPS-Tris buffer using Malvern Zetasizer (Nano-ZS, Malvern Instruments, UK).

### Bacterial bioluminescence inhibition assay and calculation of 10-min EC_50_

Constitutively luminescent recombinant bacteria *E. coli* MC1061 (pSLlux) (19) and *P. aeruginosa* DS10-129 (pDNcadlux) (18) bearing bioluminescence encoding plasmid were used. Bacterial acute bioluminescence inhibition assay (exposure time 10 min) was conducted at room-temperature on white sterile 96-well polypropylene microplates (Greiner Bio-One GmbH) following the Flash-assay protocol (ISO 21338:2010, (20)) essentially as described by Kurvet et al (21). For each compound, 5-7 sequential exponential dilutions were analysed. The following concentrations of test chemicals were used: AgNP and AgNO_3_ from 2.5 to 320 μM (0.27 to 35 mg Ag/l); PMB from 1.2 to 160 μM (1.66 to 222 mg/l); 3,5-DCP from 47.9 to 6135 μM (7.8 to 1000 mg/l). Testing was performed in MOPS-Tris environment. The 10-min EC_50_ values (the concentration of a compound reducing the bacterial bioluminescence by 50% after being in contact for 10 minutes) were calculated from the concentration *versus* 10-min inhibition curves based on nominal exposure concentrations and using the log-normal model of MS Excel macro Regtox (22).

### Evaluation of the minimal bactericidal concentration (MBC)

Since bioluminescence inhibition test do not fully reflect the viability of cells, the viability assay was performed essentially as described by Suppi et al (23). Briefly, in the end of the bioluminescence inhibition assay (see above) microplates were incubated for 1 hour at room-temperature in the dark, and 3 μl of each test suspension was pipetted onto LB agar plates to assess the viability (1-h MBC) of the cells after exposure to studied compounds. The inoculated agar plates were incubated at 30 °C for 24 h and the 1-h MBC was defined as the lowest tested concentration of the toxicant yielding no visible bacterial growth (i.e., irreversible growth inhibition) on nutrient agar.

### Assessment of bacteria-nanoparticle interaction by flow cytometry

Flow cytometry analysis was performed on the basis of protocol by Kumar et al (24). 600 pl of bacterial suspension (OD_600_~1) in 50 mM MOPS-Tris was exposed to 10 pM of AgNO_3_ or 100 μM AgNP for 0 and 10 min in polypropylene microcentrifuge tubes. Unexposed cells in 50 mM MOPS-Tris served as a control. After desired incubation time samples were vortexed and analysed using BD Accuri™ C6 (BD Biosciences). For all flow cytometry experiments data for 10 000 events was collected. The dot plots were obtained by plotting the forward scattering (FSC) intensity (X-axis, log-scale) and side scattering (SSC) intensity (Y-axis, log-scale); BD Accuri C6 software was used to analyse the results. The gating of the data as SSC-A High and SSC-A Low in individual experiments was set relatively to SSC-A of unexposed cells (control) to obtain SSC-A High of 0.2-0.5%. In addition, propidium iodide staining (in the final concentration of 5 μg/ml; exposure time 10 min) was applied to determine the viability of cells (25).

Extracellular and intracellular AgNP in exposed bacterial suspensions were discriminated using chemical procedure described by Braun et al (26). Briefly, 600 μl of bacterial suspension (OD_600_~1) in 50 mM MOPS-Tris was exposed to 10 μM of AgNO_3_ or 100 μM AgNP for 0, 10 and 30 min in microcentrifuge tubes. After exposure, cell suspension was washed (centrifuged at 6 000 g for 5 min) twice in phosphate buffered saline (PBS), incubated with two solutions (20 mM of K_3_Fe(CN)_6_ and Na2S2O3, final concentrations) for 5 min, centrifuged at 6 000 g for 5 min and resuspended in 600 μl PBS. The removal of extracellular AgNP (oxidation of cell wall-bound AgNP first to Ag^+^ ions by K_3_Fe(CN)_6_ and then followed by the complexation of Ag^+^ ions by Na_2_S_2_O_3_) was verified by colour change of AgNP from characteristic yellow to colourless. After the oxidation, the flow cytometry analysis was performed as described above.

### Assessment of interaction of studied compounds with bacterial membranes using 1-N-phenylnaphthylamine (1-NPN)

The cell wall permeabilization of *E. coli* and *P. aeruginosa* by AgNO_3_, AgNP and PMB was assayed by the cellular uptake of a hydrophobic probe 1-N-phenylnaphthylamine (1-NPN) essentially as described by Helander and Mattila-Sandholm (27). As compared to hydrophilic environments, the fluorescence of 1-NPN is significantly enhanced in hydrophobic environments (e.g. membrane lipid bilayer), rendering it a suitable dye to probe outer membrane integrity of Gram-negative bacteria (27). Briefly, 50 μl of 40 μM 1-NPN and 50 μl of tested compound in 50 mM MOPS-Tris buffer were pipetted into black microplates. 100 μl of bacterial suspension in 50 mM MOPS-Tris was dispensed into each well and the fluorescence was immediately measured (Fluoroskan Ascent FL plate luminometer; excitation/emission filters 350/460 nm). The 1-NPN cell uptake factor was calculated as a ratio between intensity of fluorescence values of the bacterial suspension incubated with and without test compounds.

### Electrochemical measurements

Bacterial cultures were grown overnight in LB broth, containing also 0.5% NaCl (Sigma-Aldrich, Munich, Germany), diluted 1:50 in fresh medium, and the incubation was continued until the OD_600_ reached 1.0. The cells were harvested by centrifugation at 4 °C for 10 min at 3000 g (Heraeus™ Megafuge™ 16R, Thermo Scientific, Germany). The pelleted cells were re-suspended in LB medium without NaCl (to avoid the formation of insoluble AgCl while exposing bacteria to silver compounds), pH 8.0, to obtain ~2×10^11^ cells ml^−1^. The concentrated cell suspensions were kept on ice until used, but not longer than 4 h.

TPP^+^ and K^+^ concentrations in the incubation media were potentiometrically monitored using selective electrodes as described previously (28, 29). In the experiment with TPP^+^, ethylenediaminetetraacetic acid (EDTA) was used to increase OM permeability of bacteria to TPP^+^ and to equilibrate TPP^+^ across the cell envelope. Since *P. aeruginosa* cells are sensitive to the incubation in EDTA-containing MOPS/Tris buffer (29), 100 mM NaPi was used instead of MOPS/Tris in the experiments with these cells. The thermostated magnetically stirred glass vessels were filled with 5 ml of 50 mM MOPS-Tris, pH 8.0. After calibration of the electrodes the concentrated cell suspension was added to obtain an OD_600_ of 1 or 2. To register potential of the electrodes we used the electrode potential-amplifying system with an ultralow-input bias current operational amplifier AD549JH (Analog Devices, Norwood, MA, USA). The data acquisition system PowerLab 8/35 (ADInstruments, Oxford, UK) was used to connect the amplifying system to a computer. The agar salt bridges were used for indirect connection of the Ag/AgCl reference electrodes (Thermo Inc.; Orion model 9001) with cell suspensions in the vessels. The measurements were performed simultaneously in 2-4 reaction vessels and three independent series of measurements were conducted.

Dissolved oxygen level in the medium was monitored by dissolved oxygen probe (Orion model 9708, Thermo Scientific) as described previously (28). At the end of every experiment solid Na_2_S_2_O_5_ (Sigma) was added to vessels to obtain a final concentration of approximately 20 mM and deplete the dissolved oxygen (0 % base line). Concentration of the dissolved oxygen in the medium before addition of cells was set as 100 %.

### Statistical analysis

All tests were performed in duplicates and in at least three independent experiments. The data were expressed as mean ± standard deviation (SD) or shown as representative figure. To define statistically significant differences, the data were analyzed with unpaired two-tailed t-test assuming equal variances at p<0.01.

## RESULTS

### Physico-chemical characterization of silver nanoparticles

AgNP formed stable dispersion in the MOPS-Tris buffer (polydispersity index of 0.31 nm) and were in nano-size of 50 nm (Table 1). The dissolution of AgNP in the test conditions was time-dependent and relatively high: 26.9 % already after 10 minutes of incubation.

To rule out the reactive oxygen species (ROS)-associated antibacterial mechanism of toxicity, the potential of AgNP to generate ROS was measured. No ROS were induced by 0.3-10 μM AgNP in abiotic conditions after 1-h incubation in 50 mM Tris-MOPS according to 2′,7′-dichlorofluorescin assay (data not shown).

**Table 1.** Characterization of silver nanoparticles (AgNP) used in the current study.

|  |  |
| --- | --- |
| Primary size, nm | 12.5±4 (16) |
| Coating | Casein, 30% |
| Hydrodynamic size in MOPS-Tris<sup>1</sup>, nm | 50 ± 0.7 |
| Polydispersity index in MOPS-Tris<sup>1</sup>, nm | 0.31 ± 0.02 |
| Surface charge of NP in MOPS-Tris<sup>1</sup>, nm | -26.6 ± 1.7 |
| Dissolution<sup>2</sup> in MOPS-Tris, % |  |
| 10 min | 26.9 ± 2.0 |
| 1 h | 31.8 ± 3.3 |
| 24 h | 66.3 ± 3.3 |
| Abiotic generation of reactive oxygen species<sup>3</sup> | No |

<sup>1</sup> Measured by dynamic light scattering (DLS); <sup>2</sup> Analyzed by atomic absorption spectroscopy from supernatants of ultracentrifuged (390 000 g\*30 min) 10 µM AgNP suspensions after 10-min, 1-h and 24-h incubation in MOPS-Tris medium at room temperature. <sup>3</sup> Reactive oxygen species (ROS) were determined using 2',7'-dichlorofluorescein diacetate as described by Aruoja et al (30) and using Mn<sub>3</sub>O<sub>4</sub> NP as positive control.

### Effect of exposure of bacteria to studied compounds on bacterial bioluminescence

The inhibition of bacterial bioluminescence was used as the toxicity endpoint describing the rapid general adverse effect of studied compounds on bacterial membranes. In bioluminescent bacteria, any disturbance of bacterial membranes leads to the decrease in the light output (inhibition of bioluminescence) that can be registered (31). As the production of light is correlating with the ATP production that proceeds in bacterial PM, the assay is suitable for the rapid analysis of the effect of chemicals on the intactness of bacterial cellular membranes. Our data showed that Ag^+^ ions and AgNP had rapid effects cell membrane-linked ATP production (Fig. 2A-D) and bacterial viability (Fig. 2E-F). As expected, in both assays *P. aeruginosa* was more susceptible to Ag^+^ ions and AgNP than *E. coli*, whereas *E. coli* was more susceptible to antibiotic PMB. 3,5-DCP that was included as the control due to its wide use as a standard in various toxicological tests (32) had similar 10-min EC_50_ and 1-h MBC values to both bacteria (Fig. 2E-F). Thus, differently from PMB and 3,5-DCP, Ag compounds were more toxic to *P. aeruginosa* compared to *E. coli*, probably becase the former cells are mostly dependent on membrane ATP-synthase (33). Thus, it can be hypothesized that Ag compunds interferred with the systems involved into ATP syntheisis.

**FIG 2.**
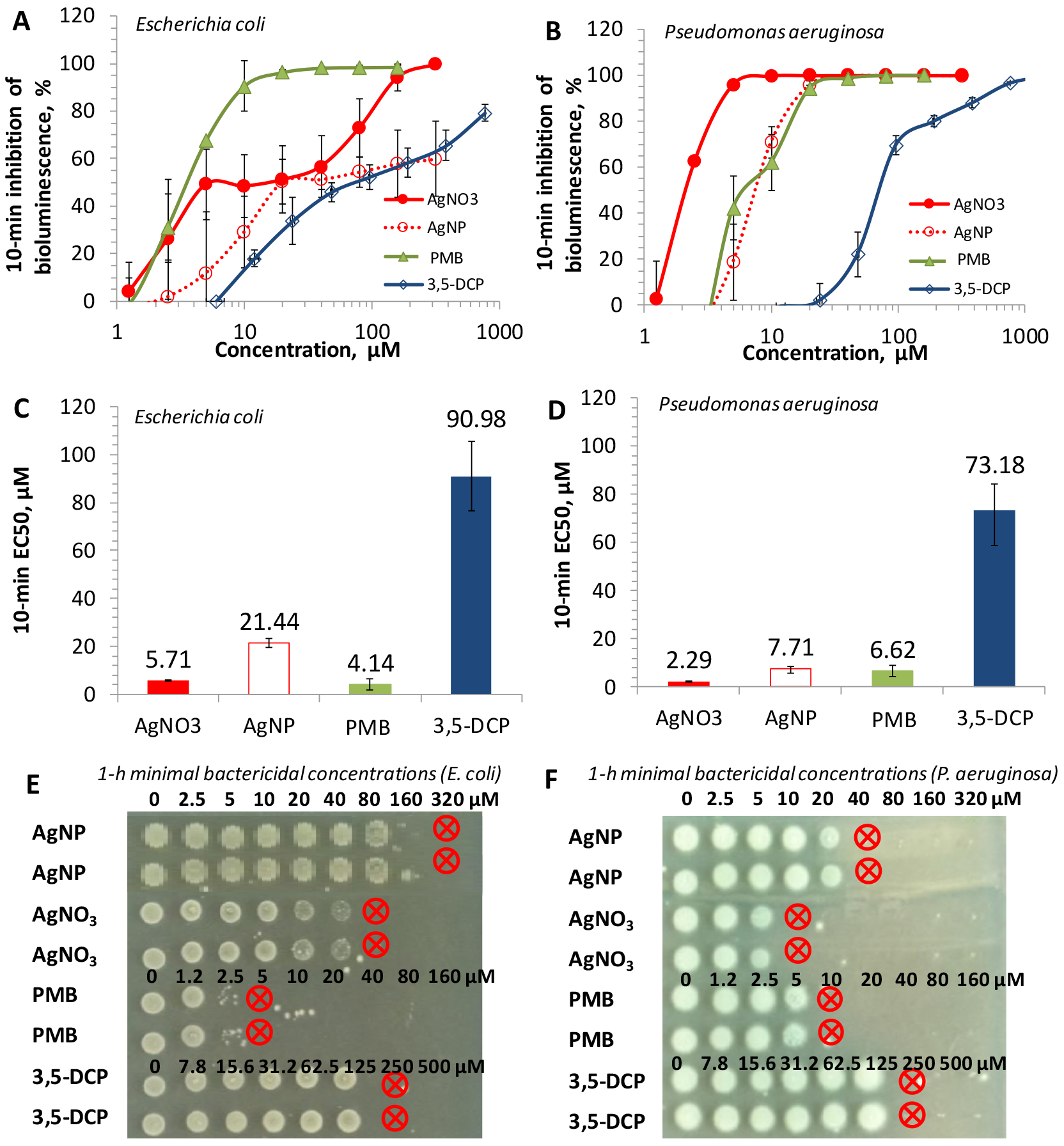
Toxicity of AgNO_3_, Ag nanoparticles (AgNP), polymyxin B (PMB) and 3,5-dichlorophenol (3,5-DCP) to bioluminescent *E. coli* and *P. aeruginosa*. **A-B:** Dose-response curves (inhibition of bioluminescence in bacteria) after 10-min exposure to Ag compounds, PMB and 3,5-DCP in MOPS-Tris buffer. **C-D:** 10-min EC_50_-s calculated from the dose-response curves presented in panels A and B, n=5; average± SD are shown. **E-F:** Minimal bactericidal concentrations of Ag compounds, PMB and 3,5-DCP after 1-h incubation (1 h MBC, pM). Red circles mark the MBC values. Data of two replicates for each compound are presented.

In viability assay, Ag compounds completely abolished bacterial growth at concentrations from 10 μM (AgNO_3_) to 40 μM (AgNP) for *P. aeruginosa* and from 80 μM (AgNO_3_) to 340 μM (AgNP) *E. coli* (Fig. 2C-D). The four-times difference between the toxicity of AgNO_3_ and AgNP evident for both bacteria was most probably at least partly due to dissolution of AgNP (26.9 % in the test medium, Table 1).

### Assessment of cell-nanoparticle interaction by flow cytometry

Bacterial suspensions were incubated either with 10 μM AgNO_3_ or 100 μM AgNP, and the intensity of the side scattering (SSC-A) was visualized and quantified by flow cytometry (Fig. 3A-B). SSC-A is an universal parameter reflecting the changes in cell granularity and is often interpreted as an indication of the internalization of NP by cells (24, 34). However, we also considered that interaction of AgNO_3_/AgNP with the cell envelope from outside can lead to the disorganization of cell membranes (9) and changes in the light scattering. Thus, both, the outside interaction of the cells with NP and actual internalization of NP by the cells can potentially lead to the increase of SSC-A. Flow cytometry analysis showed that the incubation of *P. aeruginosa* cells with 100 μM AgNP increased the intensity of SSC-A (Fig. 3A). The 33.2 % increase of SSC-A of AgNP-exposed *P. aeruginosa* cells was observed compared to unexposed cells already after 10 minutes of exposure (Fig. 3B). Remarkably, the incubation of *P. aeruginosa* cells with 10 μM AgNO_3_ did not cause the SSC-A changes, showing that AgNP interacted with *P. aeruginosa* cells in a specific manner, differently from the effects of ionic silver.

**FIG 3.**
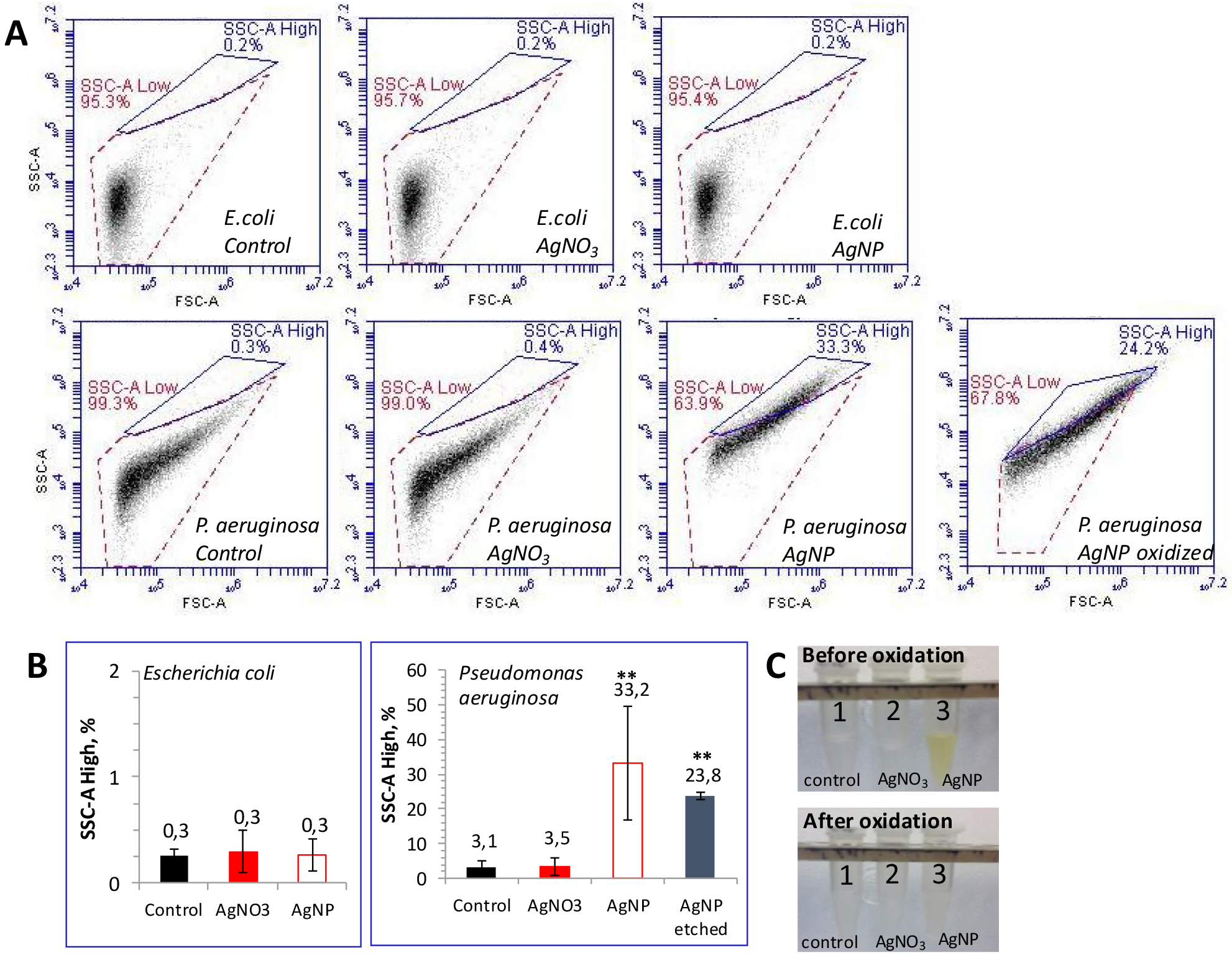
Flow cytometry analysis of *E. coli* and *P. aeruginosa* cells incubated with 10 μM AgNO_3_ or 100 μM Ag nanoparticles (AgNP). **A:** Representative figures of *E. coli and P. aeruginosa* cells showing the absence of side scatter (SSC-A) shift in case of *E. coli* (upper panels) and the presence of SSC-A shift in case of *P. aeruginosa* (lower panels) incubated with AgNP. **B:** Changes in the intensity of SSC-A calculated from the raw data shown on panel A, n=3 ± SD are shown. **C:** *P. aeruginosa* cells in MOPS-Tris buffer (control, 1) or incubated with 10 μM AgNO_3_ (2) or 100 μM AgNP (3) for 10 minutes. After the oxidation AgNP (3) lost their characteristic yellowish color.

To exclude the possibility that SSC-A shift was an artefact indicating the death of *P. aeruginosa* cells after the 10-minute exposure to AgNP, we measured the bacterial viability using fluorescein diacetate dye that stains all cells (viable+dead) (23) and propidium iodide (PI) dye that stains only dead cells (25). The viability of unexposed cells according to PI staining was over 90%. The viability of cells exposed to 100 μM AgNP and 10 μM Ag ions was over 85% for *E. coli* and over 75% for *P. aeruginosa* (data not shown). As the viability of *P. aeruginosa* cells exposed to Ag ions and AgNP was similar but the SSC-A shift was observed only for AgNP, we concluded that the shift of SSC-A was indeed observed due to effects of AgNP and not because of death of the cells or possible membrane changes in the dying.

No changes in SSC-A for *E. coli* cells was observed at any time point (Fig. 3A-B) suggesting that AgNP did not attach to the *E. coli* cell envelope and did not enter the bacterial cell or this interaction was weaker than in case of *P. aeruginosa*.

We further asked, whether the SSC-A shift in case of *P. aeruginosa-AgNP* interaction was caused by the outside adsorption of AgNP to bacterial cells walls or the actual internalization of AgNP by bacteria. To descriminate between intra- and extracellular AgNP, the method described by Braun et al (26) was used, i.e., two chemicals were utilized to remove (oxidize) NP and Ag ions from the bacterial surface. The oxidizing efficiency was verified by the visual observations, since AgNP lost their characteristic yellowish color after the oxidation (Fig. 3C). Flow cytometry analysis showed that the SSC-A shift observed for *P. aeruginosa* cells incubated with AgNP was still observed even after several washing rounds of the cells (Fig. 3A-B), suggesting that some fraction of AgNP crossed at least the OM and localized in the μM or cytosol.

### Evaluation of outer membrane integrity by 1-N-phenylnaphthylamine assay

The impact of Ag^+^ ion and AgNP on the barrier properties of the cell envelope was studied using nonpolar dye 1-N-phenylnaphthylamine (1-NPN) that fluorescesnce strongly in the lipid environment but only weakly in the hydrophilic medium (27). A cationic peptide μMB was used as a positive control due to its ability to disrupt the OM barrier and diminish the membrane potential (7). When the membrane-active compounds disturb the integrity of OM, lipophilic phase of cell envelope becomes accessible to hydrophobic agents, such as 1-NPN. Thus, the increase of 1-NPN fluorescence could be interpreted as the loss of OM integrity (27). Alternatively, the fluoresence can increase in case of the general de-energization of cells leading to the inhibition of efflux of lipophilic compounds, including 1-NPN (35).

The results of our experiments showed that membrane-permeabilizing antibiotic μMB increased the 1-NPN fluorescence approximately 3-fold in both *E. coli* and *P. aeruginosa* suspensions (Fig. 4A and B). AgNO_3_ affected the 1-NPN fluorescence in case of *E. coli* cells starting from 5 μM and AgNP starting from 40 μM, although their effect was weaker than in case of PMB. As the dissolution of AgNP was 26.9% at these conditions (Table 1), the effects of AgNP on 1-NPN fluorescence was most probably caused by Ag^+^ ions released from AgNP.

**FIG 4.**
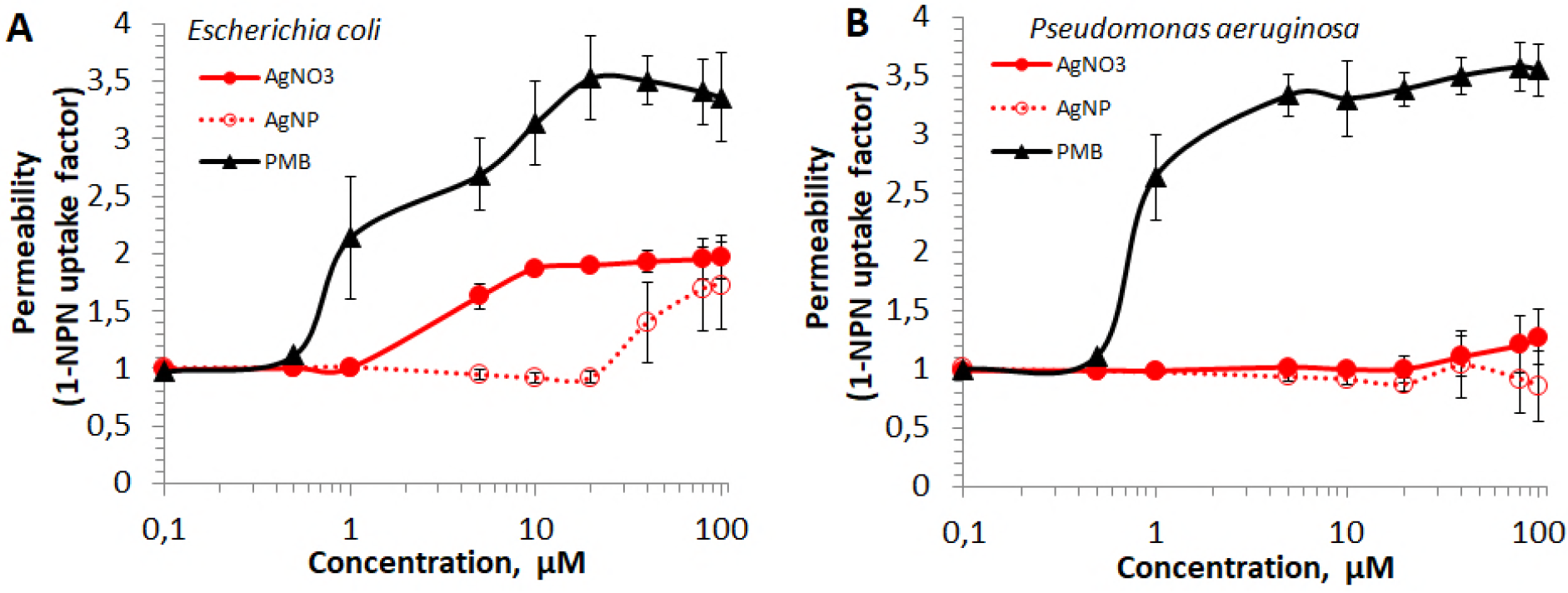
Bacterial outer membrane integrity upon exposure to AgNO_3_, Ag nanoparticles (AgNP) or polymyxin B (PMB) during 10 minutes. Fluorescence of 1-NPN was used as a permeability marker. **A:** *E. coli*, **B:** *P. aeruginosa*.

Differently from *E. coli*, no effect of Ag compounds (neither AgNO_3_, nor AgNP) was observed in the case of *P. aeruginosa* suspensions (Fig. 4B). This was in contrast to higher toxicity of AgNO_3_ and to AgNP towards *P. aeruginosa* compared to *E. coli* cells (Fig. 2) and the specific interaction of AgNP with *P. aeruginosa* AgNP cells (Fig. 3). These data suggest that the OM is not a primary target of Ag compounds. Thus, we analyzed AgNO_3_ and AgNP effects on bacteria in more details monitoring their interaction with PM and OM using TPP^+^ assay.

### Evaluation of the permeability of bacterial membranes by TPP^+^ assay

To assay effects of AgNP and Ag^+^ ions on the permeability of bacterial OM and PM, TPP^+^ accumulation inside the cells was analyzed. The OM (Fig. 1) forms a permeability barrier to lipophilic ions like TPP^+^. On the other hand, PM membrane is permeable to TPP^+^ and this indicator compound accumulates in bacterial cytosol in potential-dependent mannier. Thus, accumulation of TPP^+^ by the cells would indicate permeabilization of the OM by the Ag compounds, whereas leakage of the accumulated TPP^+^ would indicate the depolarization of PM (7). *E. coli* and *P. aeruginosa* cells with the intact OM accumulated low amounts of TPP^+^ and additions of increasing concentrations of AgNO_3_ to bacterial suspensions did not induce any additional binding of TPP^+^ to the cells, indicating that Ag compounds do not act on OM.

To induce the accumulation of TPP^+^ and to measure the effect of Ag compounds on PM, EDTA was added to the cells. EDTA removes divalent cations from the LPS layer, increasing the permeability of the OM to lipophilic compounds (7). Both bacteria accumulated considerable amounts of TPP^+^ when 0.1 mM EDTA was added to the cell suspensions (Fig. 5). 1-2.5 μM AgNO_3_ did not induce any changes in the amount of cell-accumulated TPP^+^, but 5 μM and higher concentrations of Ag^+^ ions induced release of TPP^+^ from the cells. 20 μM Ag^+^ ions induced release of the maximal amount of TPP^+^. In experiments with AgNP, 160 μM concentration was needed to depolarize the bacterial PM and induce the leakage of accumulated TPP^+^ (Fig. 5C,D).

**FIG 5.**
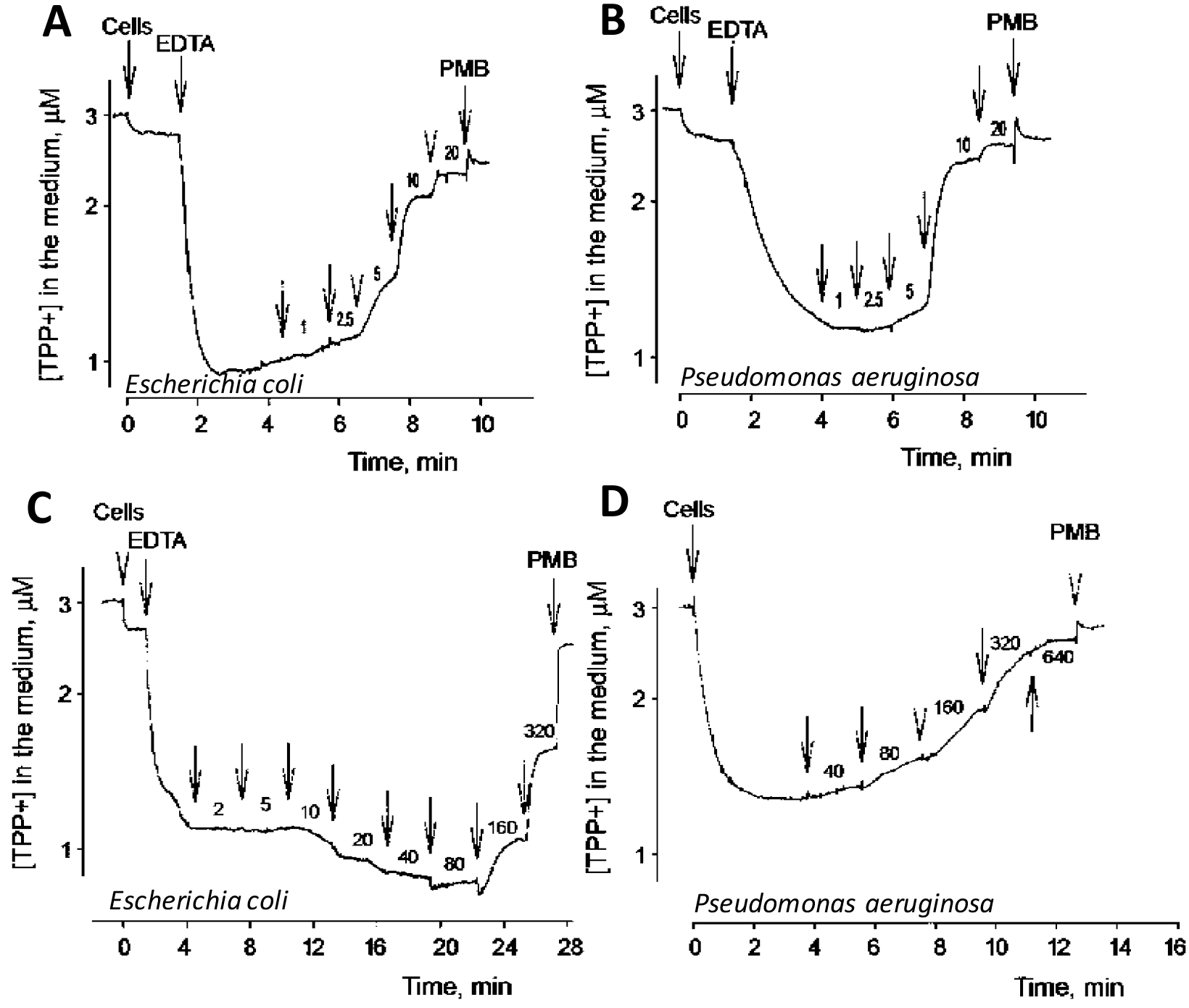
Effects of AgNO_3_ (A, B) or AgNP (C, D) on TPP^+^ accumulation by *E. coli* (A, C) and *P. aeruginosa* (B, D) cells. Experiments were performed at 37 °C in 50 mM MOPS-Tris (for *E. coli*) or 100 mM NaPi (for *P. aeruginosa*) buffers, pH 8.0. The cells were added to OD 2. Arrows (if not indicated otherwise) indicate additions of AgNO_3_ or AgNP, numbers next to the arrows indicate Ag concentrations (μM) after the last addition. EDTA was added to the final concentration of 0.1 mM, and PMB - to 100 μg/ml. In Fig 5 D 0.1 mM EDTA was added to the medium before the cells.

### Evaluation of the permeability of the bacterial plasma membrane by K^+^ leakage assay

K^+^ ions easily cross the OM through porins but the PM forms a barrier to this ion. Concentration of K^+^ ions in incubation medium was monitored to evaluate the effects of AgNP and Ag^+^ ions on barrier functions of the PM. 10 μM AgNO_3_ induced a leakage of accumulated K^+^ from the cells of both bacteria (Fig. 6). Induction of the leakage of accumulated K^+^ ions at higher Ag^+^ concentrations than leakage of TPP^+^ indicates that depolarization of the PM is most likely the reason of dissipation of K^+^ gradient. Unfortunately, it was technically not possible to register AgNP-induced K^+^ release since high concentrations AgNP affected the K^+^ electrode.

**FIG 6.**
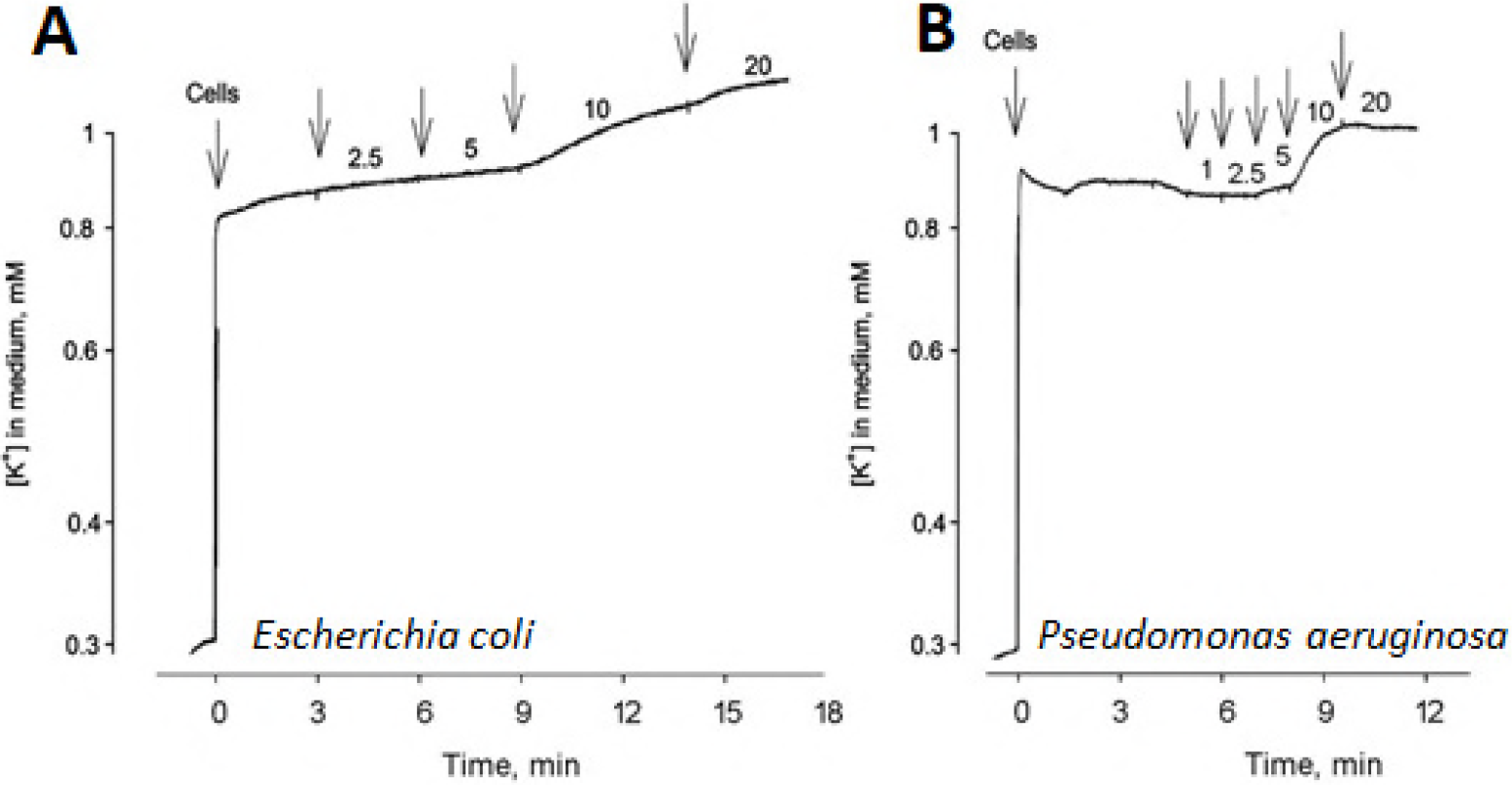
Effect of Ag^+^ on K^+^ accumulation in *E. coli* (A) and *P. aeruginosa* (B) cells. Experiments were performed at 37 °C in 50 mM MOPS-Tris buffer (for *E. coli*) or 100 mM NaPi (for *P. aeruginosa*) buffers, pH 8.0, the cells were added to OD 2. Arrows (if not indicated otherwise) indicate additions of AgNO_3_, numbers next to the arrows indicate Ag^+^ concentrations (μM) after the last addition.

### Evaluation of the permeability of bacterial plasma membrane using cell respiration assay

To understand the mechanism of Ag^+^ ion and AgNP action on bacterial PM, we analysed the respiration activity of the cells. Starting from 1 μM concentration the addition of AgNO_3_ stimulated the respiration of *E. coli* cells but in 5 minutes the activation period was followed by a considerable Ag^+^ concentration-dependent inhibition (Fig. 7A). With the increase of Ag^+^ concentration, period of the enhanced respiration activity became shorter and the effect of inhibition stronger. Period of the stimulated respiration of *E. coli* cells was considerably less expressed in the medium without glucose (data not shown). AgNP did not have any respiration stimulating activity at these conditions (Fig. 7B). In the case of *P. aeruginosa* cells, the respiration enhancing activity of AgNO_3_ was the highest at 5 μM, and the inhibitory activity of Ag^+^ was observed only at 10 μM. However, the AgNP started to stimulate respiration at 40 μM and the inhibition started at 160 μM (Fig. 7D). Such activation of respiration at low concentrations and inhibition at the higher ones is typical for uncouplers of oxidative phosphorylation, i.e., compounds permeabilizing the PM to H^+^ ions (36).

**FIG 7.**
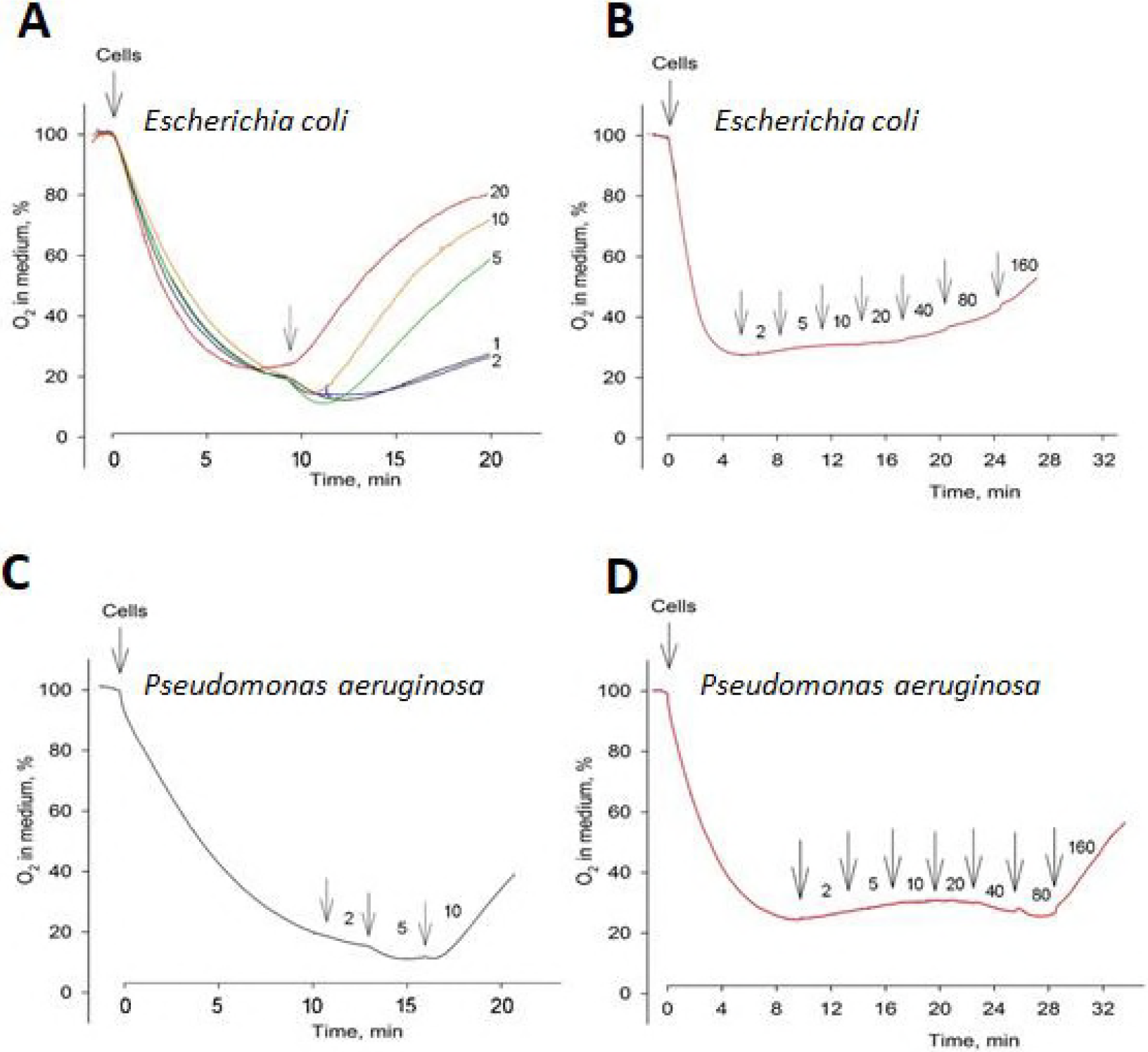
Effect of Ag^+^ (A, C) and Ag nanoparticles (B, D) on respiration of *E. coli* (A, B) and *P. aeruginosa* (C, D) cells. Experiments were performed at 37 °C in 50 mM Tris/Mops containing 0.2% of glucose (A), this medium without glucose (B), or 100 mM NaPi without glucose (C and D), all pH 8.0. The cells were added to OD 1. Arrows (if not indicated otherwise) indicate additions of AgNO_3_ or AgNP, numbers next to the arrows indicate Ag concentrations (μM) after the last addition.

In addition to *P. aeruginosa* DS10-129, *P. aeruginosa* PAO1 strain was used in the TPP^+^, K^+^ leakage and respiration assays. The results for these two bacterial strains were very similar (data not shown), suggesting that the effect of Ag is not specific to one *P. aeruginosa* strain.

### TPP^+^ assay using oxidized NP

To get direct evidences that Ag^+^ ions are responsible for the PM-damaging effects of AgNP, the experiments with the oxidized AgNP were performed. Suspensions of AgNP (final concentration 1 mM Ag) were made in KMnO_4_ solutions of concentrations from 0.05 to 0.5 mM (Fig. 8). When the concentration of the KMnO_4_ in the solution was 0.05 mM, 80 μM of AgNP was needed to induce the depolarization. However, when the concentration of KMnO_4_ was increased to 0.5 mM and AgNP were fully oxidized, only 10 μM of AgNP was needed to induce the depolarization of the PM. Thus, the efficiency of AgNP depended on the concentration of KMnO_4_ in the solution, i.e., on the oxidized fraction of Ag in the AgNP. Remarkably, the efficiency of fully oxidized AgNP (Fig. 8) was very close to the efficiency of AgNO_3_ solution (5 μM, Fig. 5). Thus, the experiments with suspensions of AgNP in KMnO_4_ solutions clearly showed, that Ag^+^ ions are responsible for the PM depolarizing effect of AgNP.

**FIG 8.**
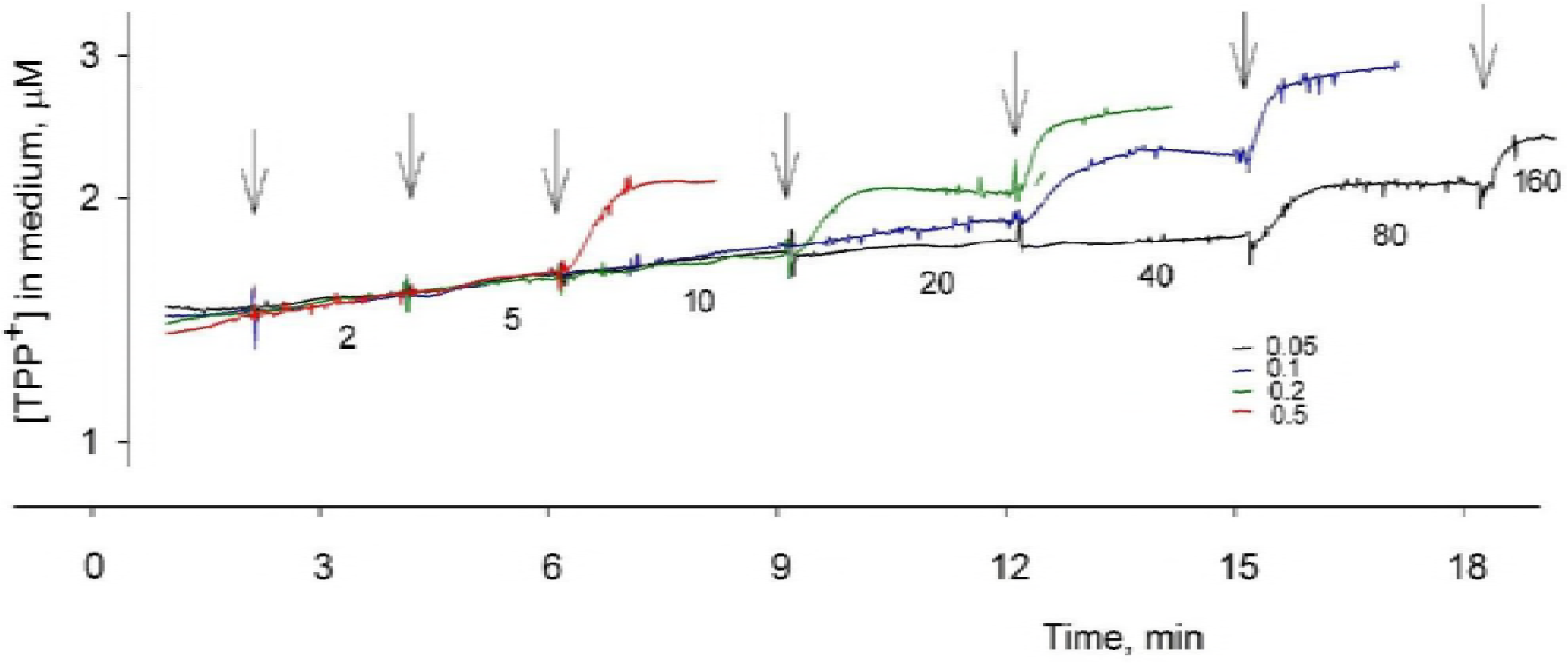
Effect of the oxidized Ag nanoparticles on TPP^+^ accumulation by *E. coli* cells. The experiments were performed at 37 °C in 50 mM Tris/Mops buffer, containing 0.1 mM EDTA, pH 8.0, when the OD of bacterial suspension was 2. Black arrows indicate the additions of AgNP containing KMnO_4_ solutions of different concentrations. Numbers under the curves between arrows indicate concentrations of Ag in the medium after the last addition of AgNP suspension in KMnO_4_ solutions.

## DISCUSSION

Discussions on the mechanisms of AgNP interaction with the bacterial cells are continuing despite of the large set of experimental data on their toxicity and possible mode of action (37, 38, 39, 40). One among the well-established toxicity mechanisms of AgNP is their adhesion to the surface of bacterial envelopes, disruption of the integrity of membranes and subsequent damage of (intra)cellular biomolecules (41). To the best of our knowledge, the exact mechanism of damage of the envelope barrier and the toxicity initiating event of AgNP are not yet defined.

In this study we showed that the PM is the main target of AgNP in *E. coli* and *P. aeruginosa*. In the bioluminescence inhibition assay (Fig. 2) we observed that *P. aeruginosa* is inherently more susceptible to Ag compounds than *E. coli*, probably because the energy production of the former cells mainly depends on PM-linked ATP-ase (33). Using TPP^+^, K^+^ and respiration assays, we confirmed that the antibacterial effects of Ag and AgNP occur through PM. Indeed, exposure of bacteria to AgNP induced a set of toxic events targeting PM: inhibition of the respiration, depolarization of the PM and the leakage of the intracellular K^+^ (Fig. 5–7). In addition, as many previous studies, we confirmed that AgNP toxicity is mediated *via* Ag ions (Fig. 8). The former was observed in bioluminescence inhibition assay (Fig. 2) in combination with dissolution studies (Table 1) and confirmed by AgNP oxidation assay that showed Ag^+^ ions dissolving from AgNP are driving the quick toxic effects of AgNP on the bacterial PM (Fig. 8). Most likely, the action of Ag^+^ ions on PM is unspecific and occurs through the binding of Ag^+^ ions with the SH-groups of various enzymes localizing on PM (12, 15). PM-related mechanism of toxicity can explain why Gram-negative bacteria including clinical multidrug-resistant strains, are in general more susceptible to AgNP then Gram-positive bacteria (40, 42). Compared to the Gram-negative bacteria, the plasma membrane of Gram-positive bacteria is less accessible to the NP due to thick peptidoglycan layer. Thus, combination of AgNP with traditional antibiotics offers crucial therapeutic strategy on fighting multidrug-resistant Gram-negative bacteria.

Surprisingly, we did not observe any effects of Ag compounds on the bacterial outer membrane (Fig. 4–5). Although AgNP clearly attached to *P. aeruginosa* cells (Fig. 3), this did not lead to the permeabilization of OM (Fig. 4–5). Several studies have previously shown structural changes in the surface of bacteria after exposure to AgNP and proposed that physical interaction with AgNP was the primary cause of cell death (13, 43). For example, Ramalingam et al (43) observed altered cellular morphology and diminished cellular integrity after exposure of *E. coli* and *P. aeruginosa* cells to AgNP and suggested that AgNP damage the cell membrane *via* binding to the cell surface proteins and lipopolysaccharides. Our study indicates that the observed morphological changes on bacterial surface could be rather the consequences of the action of Ag^+^ ions than the primary mechanism of AgNP cytotoxicity. The uptake of Ag^+^ ions leads to the formation of characteristic irregular pits on the surface of bacteria (44) that can be misleadingly interpreted as the toxicity initiating event of AgNP. Furthermore, depolarization of the PM induced by Ag^+^ ions can destabilize the OM and trigger the morphological changes of microbial envelope.

Since Ag^+^ ions dissolved from AgNP reached bacterial PM without causing a significant OM damage, these data support the idea that Ag^+^ transport occurs *via* cation selective OM porins (45, 46). Previous studies showed that *E. coli* mutant strains deficient in OmpF or OmpC porins were more resistant to Ag^+^ ions (9) and to AgNP (11) as compared to the wild type strain. In addition, *ompF* gene was down-regulated in response to Ag^+^ ions and AgNP (10) suggesting that porins are crucial in antibacterial effects of Ag^+^ ions and AgNP. On the basis of our study and the previous observations, we suggest that Ag^+^ ions dissolved from AgNP enter the bacterial periplasm through porin channels and interact directly with the PM. Previous study found that Omp proteins were among the molecules tightly bound to AgNP after their incubation with the cell free extract of *E. coli*, implying that in addition to Ag^+^ ions, AgNP under 10 nm can also pass through the porin channels to PM (40, 47). This can explain intracellular localization of AgNP in *P. aeruginosa* cells observed in this study. However, we do not have the explicit answer, why AgNP were associated and internalized by *P. aeruginosa* but not *E. coli* cells (Fig. 3). One possible explanation could be the bigger porin channels in OM of *P. aeruginosa*. It is known that channels formed by protein F in *P. aeruginosa* OM are much larger than *E. coli* porin channels (48). In addition, the OM of *P. aeruginosa* cells possess high number of different porins (49). Since AgNP are oxidized on the surface of bacteria, creating the large pools of toxic Ag ions (4, 10) and reducing the size of AgNP, the number and size of porins available for the transport of Ag^+^ ions and small AgNP can be one of the determinants of the toxicity of AgNP to different bacteria.

Finally, the assays used in our study showed the short-term (0 - 1 h) dissolution-dependent antimicrobial mechanisms of Ag compounds to both bacteria. We observed that the short-term toxicity of Collargol to *E. coli* and *P. aeruginosa* differed only several times (Fig. 2) in this study, while the long-term toxicity of Collargol was almost two orders of magnitude different for these two bacteria as shown in our previous article (4). Thus, the “quick” dissolution-dependent effects of AgNP might be universal but the time-dependent effects seem to be specific to different bacteria. For example, the long-term effects may be related to the capacity of bacteria to oxidase adsorbed AgNP to Ag^+^ ions or/and the efficiency of Ag^+^ efflux. For instance, it was shown that *E. coli* with the more active efflux transporters of Ag^+^ ions (e.g., SilCFBA) are more resistant to Ag (50). The full oxidation of adsorbed AgNP to Ag^+^ ions can take hours. The results of this slow oxidation process are not registered by our electrochemical analysis but can be reflected in the results of the long-term endpoints, i.e. the cell ability to grow. Thus, longer incubation times and bacterial growth-related endpoints might be more relevant and useful to reveal the whole toxic potential of NP *per se* compared to the effect of dissolved ionic fraction.

## CONCLUSIONS

We demonstrated that Ag^+^ ions and AgNP induced a set of toxic effects targeting PM of *E. coli* and *P. aeruginosa*, including inhibition of the respiration, depolarization, leakage of the intracellular K^+^ and the decrease of ATP content in the cells. These fast effects of AgNP depended on the release of Ag ions. The role of Ag compounds on bacterial OM and the role of particles *per se* was negligible in our short-term (0-1 h) tests.

## ACKNOWLEDGEMENTS

This work was supported by Estonian Research Council grants IUT23-5 and PUT1015 and by Research Council of Lithuania, funding grant No. MIP-040/2015. The funders had no role in study design, data collection and interpretation, or the decision to submit the work for publication.

## REFERENCES

(1) Nowack B, Krug HF, Height M. 2011. 120 years of nanosilver history: implications for policy makers. Environ Sci Technol 45:1177–1183.

(2) Djurišić AB, Leung YH, Ng AM, Xu XY, Lee PK, Degger N, Wu RS. 2015. Toxicity of metal oxide nanoparticles: mechanisms, characterization, and avoiding experimental artefacts. Small 11:26–44.

(3) Ouay B, Stellacci F. 2015. Antibacterial activity of silver nanoparticles: a surface science insight. Nano Today 10:339–354.

(4) Bondarenko O, Ivask A, Käkinen A, Kurvet I, Kahru A. 2013. Particle-cell contact enhances antibacterial activity of silver nanoparticles. PLoS One 8:e64060.

(5) Silhavy TJ, Kahne D, Walker S. The bacterial cell envelope. 2010. Cold Spring Harb Perspect Biol 2:a000414.

(6) Amro NA, Kotra LP, Wadu-Mesthrige K, Bulychev A, Mobashery S, Liu G. 2000. High-resolution atomic force microscopy studies of the Escherichia coli outer membrane: structural basis for permeability. Langmuir 16:2789–2796.

(7) Daugelavičius R, Bakiené E, Bamford DH. 2000. Stages of Polymyxin B interaction with the Escherichia coli cell envelope. Antimicrob Agents Chemother 44:2969–2978.

(8) Holt KB, Bard AJ. 2005. Interaction of silver(I) ions with the respiratory chain of Escherichia coli: an electrochemical and scanning electrochemical microscopy study of the antimicrobial mechanism of micromolar Ag^+^. Biochemistry 44:13214–13223.

(9) Li XZ, Nikaido H, Williams KE. 1997. Silver-resistant mutants of Escherichia coli display active efflux of Ag^+^ and are deficient in porins. J Bacteriol 179:6127–6132.

(10) McQuillan JS, Infante GH, Stokes E, Shaw AM. 2012. Silver nanoparticle enhanced silver ion stress response in Escherichia coli K12. Nanotoxicology 6: 857–866.

(11) Radzig MA, Nadtochenko VA, Koksharova OA, Kiwi J, Lipasova VA, Khmel IA. 2013. Antibacterial effects of silver nanoparticles on gram-negative bacteria: influence on the growth and biofilms formation, mechanisms of action. Colloids Surf B Biointerfaces 102:300–306.

(12) Dibrov P, Dzioba J, Gosink KK, Häse CC. 2002. Chemiosmotic mechanism of antimicrobial activity of Ag(+) in Vibrio cholerae. Antimicrob Agents Chemother 46:2668–2670.

(13) Sondi I, Salopek-Sondi B. 2004. Silver nanoparticles as antimicrobial agent: a case study on E. coli as a model for Gram-negative bacteria. J Colloid Interface Sci 275:177–182.

(14) Lok CN, Ho CM, Chen R, He QY, Yu WY, Sun H, Tam PK, Chiu JF, Che CM. 2006. Proteomic analysis of the mode of antibacterial action of silver nanoparticles. J Proteome Res 5:916–924.

(15) Li WR, Xie XB, Shi QS, Zeng HY, Ou-Yang YS, Chen YB. 2010. Antibacterial activity and mechanism of silver nanoparticles on Escherichia coli. Appl Microbiol Biotechnol 85:1115–1122.

(16) Blinova I, Niskanen J, Kajankari P, Kanarbik L, Käkinen A, Tenhu H, Penttinen OP, Kahru A. 2013. Toxicity of two types of silver nanoparticles to aquatic crustaceans Daphnia magna and Thamnocephalus platyurus. Environ Sci Pollut Res Int 20:3456–63.

(17) ISO/IEC, 2017. General requirements for the competence of testing and calibration laboratories. International Standard, reference number ISO/IEC 17025:2017(E). Switzerland.

(18) Bondarenko O, Rahman PKSM, Rahman TJ, Kahru A, Ivask A. 2010. Effects of rhamnolipids from Pseudomonas aeruginosa DS10-129 on luminescent bacteria: toxicity and modulation of cadmium bioavailability. Microb Ecol 59:588–600.

(19) Ivask A, Rölova T, Kahru A. 2009. A suite of recombinant luminescent bacterial strains for the quantification of bioavailable heavy metals and toxicity testing. BMC Biotech 9:41.

(20) ISO, 2010. Water quality - Kinetic determination of the inhibitory effects of sediment, other solids and coloured samples on the light emission of Vibrio fischeri (kinetic luminescent bacteria test). International Standard, reference number ISO 21338:2010(E). International Organization for Standardization, Geneva, Switzerland.

(21) Kurvet I, Ivask A, Bondarenko O, Sihtmäe M, Kahru A. 2011. LuxCDABE-transformed constitutively bioluminescent Escherichia coli for toxicity screening: comparison with naturally luminous Vibrio fischeri. Sensors 11:7865–7878.

(22) Vindimian E. 2009. http://www.normalesup.org/~vindimian/enindex.html.

(23) Suppi S, Kasemets K, Ivask A, Künnis-Beres K, Sihtmäe M, Kurvet I, Aruoja V, Kahru, A. 2015. A novel method for comparison of biocidal properties of nanomaterials to bacteria, yeasts and algae. J Hazard Mater 286:75–84.

(24) Kumar A, Pandey AK, Singh SS, Shanker R, Dhawan A. 2011. A flow cytometric method to assess nanoparticle uptake in bacteria. Cytometry A 79:707–12.

(25) Stiefel P, Schmidt-Emrich S, Maniura-Weber K, Ren Q. 2015. Critical aspects of using bacterial cell viability assays with the fluorophores SYTO9 and propidium iodide. BMC Microbiol 15:36.

(26) Braun GB, Friman T, Pang HB, Pallaoro A, Hurtado de Mendoza T, Willmore AM, Kotamraju VR, Mann AP, She ZG, Sugahara KN, Reich NO, Teesalu T, Ruoslahti E. 2014. Etchable plasmonic nanoparticle probes to image and quantify cellular internalization. Nat Mater 13:904–911.

(27) Helander IM, Mattila-Sandholm T. 2000. Fluorometric assessment of Gram-negative bacterial permeabilization. J Appl Microbiol 88:213–219.

(28) Daugelavičius R, Gaidelyté A, Cvirkaité-Krupovič V, Bamford DH. 2007. On-line monitoring of changes in host cell physiology during the one-step growth cycle of Bacillus phage Bam35. J Microbiol Methods 69:174–179.

(29) Daugelavičius R, Buivydas A, Senčilo A, Bamford DH. 2010. Assessment of the activity of RND-type multidrug efflux pumps in Pseudomonas aeruginosa using tetraphenylphosphonium ions. Int J Antimicrob Agents 36:234–238.

(30) Aruoja V, Pokhrel S, Sihtmäe M, Mortimer M, Mädler L, Kahru A. 2015. Toxicity of 12 metal based nanoparticles to algae, bacteria and protozoa. Environ Sci Nano 2:630–644.

(31) Bulich, A.A. 1982. A practical and reliable method for monitoring the toxicity of aquatic samples. Process Biochem 17:45–47.

(32) Elnabarawy MT, Robideau RR, Beach SA. 1988. Comparison of three rapid toxicity test procedures: Microtox®, Polytox®, and activated sludge respiration inhibition. Toxic. Assess 3:361–370.

(33) Temple LM, Sage AE, Schweizer HP, Phibbs PV. 1998. Carbohydrate Catabolism in Pseudomonas aeruginosa. In: Montie T.C. (eds) Pseudomonas. Biotechnology Handbooks, vol 10. Springer, Boston, MA

(34) Ivask A, Visnapuu M, Vallotton P, Marzouk ER, Lombi E, Voelcker NH. 2016. Quantitative multimodal analyses of silver nanoparticle-cell interactions: Implications for cytotoxicity. NanoImpact 1:29–38.

(35) Ocaktan A, Yoneyama H, Nakae T. 1997. Use of fluorescence probes to monitor function of the subunit proteins of the MexA-MexB-oprM drug extrusion machinery in Pseudomonas aeruginosa. J Biol Chem 272:21964–21969.

(36) Nicholls D, Ferguson SJ. 2013. Bioenergetics. 4th ed. San Diego: Academic Press, eBook ISBN: 9780123884312.

(37) Ivask A, Juganson K, Bondarenko O, Mortimer M, Aruoja V, Kasemets K, Blinova I, Heinlaan M, Slaveykova V, Kahru A. 2014. Mechanisms of toxic action of Ag, ZnO and CuO nanoparticles to selected ecotoxicological test organisms and mammalian cells in vitro: a comparative review. Nanotoxicology 8 Suppl 1:57–71.

(38) Juganson, K., Ivask, A., Blinova, I., Mortimer, M., and Kahru A. 2015. NanoE-Tox: New and in-depth database concerning ecotoxicity of nanomaterials. Beilstein J Nanotechnol 6, 1788–1804.

(39) Bondarenko OM, Heinlaan M, Sihtmäe M, Ivask A, Kurvet I, Joonas E, Jemec A, Mannerström M, Heinonen T, Rekulapelly R, Singh S, Zou J, Pyykkö I, Drobne D, Kahru A. 2016. Multilaboratory evaluation of 15 bioassays for (eco)toxicity screening and hazard ranking of engineered nanomaterials: FP7 project NANOVALID. Nanotoxicology 10:1229–42.

(40) Slavin YN, Asnis J, Häfeli UO, Bach H. 2017. Metal nanoparticles: understanding the mechanisms behind antibacterial activity. J Nanobiotechnology 15:65.

(41) Dakal TC, Kumar A, Majumdar RS, Yadav V. 2016. Mechanistic Basis of Antimicrobial Actions of Silver Nanoparticles. Front Microbiol 7:1831. eCollection 2016.

(42) Cavassin ED, de Figueiredo LF, Otoch JP, Seckler MM, de Oliveira RA, Franco FF, Marangoni VS, Zucolotto V, Levin AS, Costa SF. 2015. Comparison of methods to detect the in vitro activity of silver nanoparticles (AgNP) against multidrug resistant bacteria. J Nanobiotechnology 13: 64.

(43) Ramalingam B, Parandhaman T, Das SK. 2016. Antibacterial Effects of Biosynthesized Silver Nanoparticles on Surface Ultrastructure and Nanomechanical Properties of Gram-Negative Bacteria viz. Escherichia coli and Pseudomonas aeruginosa. ACS Appl Mater Interfaces 8:4963–76.

(44) Pal S, Tak YK, Song JM. 2007. Does the antibacterial activity of silver nanoparticles depend on the shape of the nanoparticle? A study of the Gram-negative bacterium Escherichia coli. Appl Environ Microbiol 73:1712–20.

(45) Lemire JA, Harrison JJ, Turner RJ. 2013. Antimicrobial activity of metals: mechanisms, molecular targets and applications. Nat Rev Microbiol 11:371–384.

(46) McQuillan JS, Shaw AM. 2014. Differential gene regulation in the Ag nanoparticle and Ag(+)-induced silver stress response in Escherichia coli: a full transcriptomic profile. Nanotoxicology Suppl 1:177–184.

(47) Wigginton NS, de Titta A, Piccapietra F, Dobias J, Nesatyy VJ, Suter MJF, Bernier-Latmani R. 2010. Binding of silver nanoparticles to bacterial proteins depends on surface modifications and inhibits enzymatic activity. Environ Sci Technol 44:2163–2168.

(48) Nikaido H, Nikaido, K, and Harayama. S. 1991. Identification and characterization of porins in Pseudomonas aeruginosa. J Biol Chem 266:770–779.

(49) Tamber S, Maier E, Benz R, Hancock RE. 2007. Characterization of OpdH, a Pseudomonas aeruginosa porin involved in the uptake of tricarboxylates. J Bacteriol 189:929–39.

(50) Randall CP, Gupta A, Jackson N, Busse D, O’Neill AJ. 2015. Silver resistance in Gram-negative bacteria: a dissection of endogenous and exogenous mechanisms. J Antimicrob Chemother 70:1037–46.

